# Within- and between-subject reproducibility and variability in multi-modal, longitudinal brain networks

**DOI:** 10.1101/2022.05.03.490544

**Authors:** Johan Nakuci, Nick Wasylyshyn, Matthew Cieslak, James C. Elliot, Kanika Bansal, Barry Giesbrecht, Scott T. Grafton, Jean M. Vettel, Javier O. Garcia, Sarah F. Muldoon

## Abstract

Network analysis provides new and important insights into the function of complex systems such as the brain by examining structural and functional networks constructed from diffusion Magnetic Resonance Imaging (dMRI), functional MRI (fMRI) and Electro/Magnetoencephalography (E/MEG) data. Although network models can shed light on cognition and pathology, questions remain regarding the importance of these findings, due in part to the reproducibility of the core measurements and subsequent modeling strategies. In order to ensure that results are reproducible, we need a better understanding of within- and between-subject variability over long periods of time. Here, we analyze a longitudinal, 8 session, multi-modal (dMRI, and simultaneous EEG-fMRI), and multiple task imaging data set. We first investigate the reproducibility of individual brain connections and network measures and find that across all modalities, within-subject reproducibility is higher than between-subject reproducibility, reaffirming the ability to detect individual differences in network structure in both structural and functional human brain networks. We see high variability in the reproducibility of pairwise connections between brain regions, but observe that in EEG-derived networks, during both rest and task, alpha-band connectivity is consistently more reproducible than networks derived from other frequency bands. Further, reproducible connections correspond to strong connections. Structural networks show a higher reliability in network statistics than functional networks, and certain measures such as synchronizability and eigenvector centrality are consistently less reliable than other network measures across all modalities. Finally, we find that structural dMRI networks outperform functional networks in their ability to identify individuals using a fingerprinting analysis. Our results highlight that functional networks likely reflect state-dependent variability not present in structural networks, and that the analysis of either structural or functional networks to study individual differences should depend on whether or not one wants to take into account state dependencies of the observed networks.

## 1 Introduction

The introduction of network theory to neuroscience has increased our understanding of the brain’s functional and structural organization. This powerful tool has given new insights into how higher order brain functions arise (Bassett and Sporns, 2017; Park and Friston, 2013) and how changes can lead to pathology (Fornito et al., 2015). However, questions have been raised regarding the reliability of brain network properties given the effects of noise in the signal, particularly in fMRI (Laumann et al., 2016; Power et al., 2018, 2012). Still, despite the presence of noise, brain networks have been found to exhibit consistent properties over time among individual network connections and in higher order properties, such as the clustering coefficient, characteristic path length, and assortativity, for structural connectivity as measured with dMRI (Bassett et al., 2011; Bonilha et al., 2015; Buchanan et al., 2014; Bürgel et al., 2006; Malykhin et al., 2008), fMRI (Amunts et al., 2000; Braun et al., 2012; Deuker et al., 2009; Du et al., 2015; Elliott et al., 2019; Gordon et al., 2017; Gratton et al., 2018; Laumann et al., 2015; Mangin et al., 2004; Noble et al., 2017, 2019; Pannunzi et al., 2017; Rypma and D’Esposito, 1999; Shah et al., 2016) and EEG/MEG (Deuker et al., 2009; Hardmeier et al., 2014; Kuntzelman and Miskovic, 2017). Unfortunately, most studies thus far have been limited to the analysis of a single imaging modality and/or few scanning sessions, raising questions about how reliable these properties are over longer times and across modalities.

While it is clear that there is some level of reliability in network properties within an individual over time, it is also important to understand how the state of the brain (e.g., resting wakefulness versus active task situations (Fox et al., 2005)), and the neural methodology (e.g., fMRI versus EEG) contributes to this reliability across multiple days. The “resting” brain (e.g., default mode network) is a state that has been shown to be metabolically demanding (Raichle et al., 2001) and associated not only with task performance (e.g., Tian et al., 2012) but also disease (e.g., Sorg et al., 2007), very much similar to task-related activity; however, the “resting” brain is fundamentally different from task-related activity, as engagement in a task requires precise recruitment of and coordination between regions of the brain (Fox et al., 2005). Also, in a field with a variety of diverse methodologies (e.g., fMRI, EEG, MEG, PET, etc), neuroscience researchers draw conclusions from methods that are measuring fundamentally different neural properties. For example, fMRI is an indirect measurement of neural activity, as it measures oxygenation and neural activity is inferred. Whereas EEG, a “direct” measurement, is measured on the scalp and filtered by a variety of tissues and bone separating the scalp from the brain. In terms of reliability, experimental design and task demands have shown to contribute to reliability in fMRI (Bennett and Miller, 2013, 2010), and EEG suffers from a large variety of factors that could impact reliability as well (McEvoy et al., 2000). However, there is no study, to our knowledge, that has measured reliability of network structure derived from both fMRI and EEG data collected at the same time over many sessions.

In addition to studying reliability within an individual over time, one can also ask about how network properties differ between individuals. Indeed, recent work has shown that brain networks can provide insight into the unique features associated with a person (Bansal et al., 2018b, 2018a; Gordon et al., 2017; Seitzman et al., 2019). A giant leap toward the goal of understanding differences in brain networks was made with the finding that functional brain activity has unique features that can identify a person in a group, similar to a fingerprint (Finn et al., 2015). This fingerprinting property has also been found in structural connectomes (Powell et al., 2018; Yeh et al., 2016). Fingerprinting is important because it allows neuroimaging analyses to focus on the individual and not only on group-level differences (Finn et al., 2015).

To further understand reliability in brain networks over time, across different states, and across modalities, we quantified within- and between-subject reliability in a rich longitudinal and multi-modal dataset consisting of dMRI and simultaneous EEG-fMRI recording during resting-state and multiple tasks. Importantly, the data set studied here was part of a larger study examining naturalistic sleep variability in individuals (Thurman et al., 2018). Here, we do not focus on the effects of variation in sleep pressure, but instead note that due to the study design, subjects varied in the amount of sleep pressure they experienced during each imaging session, presumably augmenting variability within- and between-subjects’ functional brain network over time. We examine both structural and functional brain networks in this data set to study reliability of individual connections and higher order network statistics. To create structural networks, dMRI imaging was used to perform tractography and network connections were defined as the density of streamlines between brain regions. fMRI networks were constructed using the Pearson-Product Correlation to quantify the magnitude of the statistical relationship in the BOLD signal between brain regions. For EEG, the time-series signal from each sensor was first separated into traditional frequency bands of δ (1-3 Hz), θ (4-7 Hz), α (8-13 Hz), β (14-30 Hz) and γ (30-60 Hz), and functional connectivity was calculated using the debiased-weighted Phase-Lag Index (dwPLI) which quantifies phase synchronization between sensors based on the consistency of the lag between the instantaneous phases of two sensors (Vinck et al., 2011).

In the current work we evaluate: 1) which brain connections and network measures are most reliable within- and between-individuals; 2) how reliability varies across state and modality; and 3) how the different imaging modalities, dMRI, fMRI, and EEG, perform in a fingerprinting analysis to identify an individual.

## 2 Material and methods

### 2.1 Participants

The University of California, Santa Barbara (UCSB) Human Subjects Committee (#16– 0154) and Army Research Laboratory Human Research Protections Office (#14–098) approved all procedures, and all participants provided informed written consent. Research was conducted in accordance with the declarations of Helsinki. The data presented in this manuscript represent a subset of data collected as part of a large-scale, longitudinal experimental that collected bi-weekly structural and functional brain data. A full description of the study can be found in (Thurman et al., 2018). Here we analyze data from 27 healthy participants who were recruited by word of mouth and local advertisements. Note that by study design, participants were excluded from the multi-session segment of the study if they did not experience sleep variability. Data is accessible upon request as far as allowed by the security policy and guidelines established with the ethics committee of the US Army Research Laboratory Human Research Protection Program.

### 2.2 Data Description

Over the course of 16 weeks, subjects were asked to complete 8 recording sessions involving dMRI and simultaneous EEG-fMRI. For each session, simultaneous EEG-fMRI recording consisted of a 5-minute resting state and 10 tasks with varying levels of cognitive demand; specifically:

Dot Probe Task (Dot) (Sipos et al., 2014);
Dynamic Attention Task (DYN 1-4) with four repetitions of the same task (Yantis et al., 2002);
Modular Math (MOD) (Mattarella-Micke et al., 2011);
Psychomotor Vigilance Task (PVT) (Loh et al., 2004), and;
Visual Working Memory (VWM 1-3) with three repetitions of the same task (Luck and Vogel, 1997).

Table 1 shows the average number of subjects and sessions for each imaging modality. When analyzing the EEG-fMRI data, we analyzed only six sessions of data. This was done in order to make a trade-off between maximizing the number of subjects and number of sessions, since not all subjects participated in all 8 sessions. Detailed information on the number of subjects and sessions for functional data can be found in Tables 2 and 3. Lastly, for the fingerprinting analysis using dMRI data, we used 25 subjects, all of which had an equal number of sessions (8 sessions). For the fingerprinting analysis using fMRI data, 15 subjects were included with all 6 sessions of resting-state and task recordings, and for the EEG data, we used 26 subjects with resting-state and all tasks over 6 sessions.

**Table 1.** Average number of subjects and sessions per imaging modality

| Imaging Modality | Subjects | Sessions |
| --- | --- | --- |
| dMRI | 25 | 8 |
| fMRI | 23.1 | 6 |
| EEG | 27 | 6 |

**Table 2.** Number of subjects and sessions for each task per EEG and fMRI

| Task | EEG Subjects | fMRI Subjects | Sessions |
| --- | --- | --- | --- |
| Resting-State | 27 | 26 | 6 |
| DOT | 27 | 25 | 6 |
| DYN-1 | 27 | 27 | 6 |
| DYN-2 | 27 | 27 | 6 |
| DYN-3 | 27 | 26 | 6 |
| DYN-4 | 27 | 20 | 6 |
| MOD | 27 | 17 | 6 |
| PVT | 27 | 19 | 6 |
| VWM-1 | 27 | 27 | 6 |
| VWM-2 | 27 | 23 | 6 |
| VWM-3 | 27 | 17 | 6 |

**Table 3.** Number of subjects and sessions for fingerprinting

| Fingerprinting | Subjects | Sessions | Tasks Included |
| --- | --- | --- | --- |
| dMRI | 25 | 8 | n/a |
| fMRI | 15 | 6 | Yes |
| EEG | 27 | 6 | Yes |

### 2.3 fMRI Acquisition and Preprocessing

Functional neuroimaging data were acquired on a 3T Siemens Prisma MRI using an echo-planar imaging (EPI) sequence (3mm slice thickness, 64 coronal slices, field of view (FoV)=192 × 192 mm, repetition time (TR)=910 ms, echo time (TE)=32 ms, flip angle=52°, and voxel size: 3 × 3 × 3 mm). For repeated scans, a T1-weighted structural image was also acquired using a high-resolution magnetization prepared rapid acquisition gradient echo (MPRAGE) sequence (TR= 2500 ms, TE=2.22 ms, and FoV= 241 × 241 mm with a spatial resolution of .9 × .9 × .9 mm), for use in coregistration and normalization.

fMRI BOLD images were preprocessed using Advanced Normalization Tools (ANTs) (Avants et al., 2009). Physiological artifacts including respiration and cardiac cycle effects were corrected using the retrospective correction of physiological motion effects method, RETROICOR (Glover et al., 2000), implemented in MEAP v1.5 (Cieslak et al., 2018). Head motion was estimated using antsMotionCorr, and the motion correction was completed as follows: (1) An unbiased BOLD template was created within each session by averaging the motion-corrected BOLD time series from each run. (2) The BOLD templates were coregistered to the corresponding T1-weighted high resolution structural images, collected in each session. (3) Each session was spatially normalized to a custom study-specific multi-modal template which included T1-weighted, T2-weighted and GFA images from twenty-four quasi-randomly selected participants chosen to match the study population. (4) The template was then affine-transformed to the coordinate space of the MNI152 Asymmetric template. (5) Finally, the fMRI volumes were transformed using the estimated head motion correction, BOLD template coregistration, BOLD-to-T1w coregistration and spatial normalization into MNI space using a single Hamming weighted sinc interpolation. After these transformations, the final step in the preprocessing was to extract time-series from fMRI scans for functional connectivity analyses. Two atlases were used to reduce the 3D volume data into 221 nodal time series data: (1) the cortical Schaefer 200 atlas (Schaefer et al., 2018) which was derived from intrinsic functional connectivity in resting state fMRI and (2) 21 subcortical regions from the Harvard-Oxford atlas based on anatomical boundaries (Makris et al., 2006). As the atlases are in MNI coordinate space, voxels within each labelled region of the atlases were simply averaged, and time series were extracted for the following connectivity analyses.

To assess functional connectivity among ROIs, mean regional time-courses were extracted and standardized using the nilearn package (Abraham et al., 2014) in Python 2.7, and confound regression was then conducted. In particular, the time series for each region was detrended by regressing the time series on the mean as well as both linear and quadratic trends. There were a total of 16 confound regressors, which included: head motion, global signal, white matter, cerebrospinal fluid and derivatives, quadratics and squared derivatives. This functional connectivity preprocessing pipeline was selected based on conclusions from prior work that examined performance across multiple commonly used preprocessing pipelines for mitigating motion artifact in functional BOLD connectivity analyses (Ciric et al., 2017; Lydon-Staley et al., 2018).

To construct the fMRI networks, the signal from all voxels within a brain region were averaged, and the Pearson Product Correlation (*R*) between two brain regions was calculated as

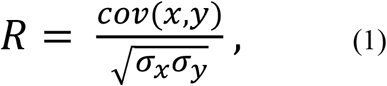

where *x* and *y* represent the time-series data from two different regions and σ is the variance of the time series. To account for negative correlations, the absolute value of the correlations was used to construct weighted functional connectivity matrices.

### 2.4 EEG Acquisition and Preprocessing

Continuous EEG recordings were captured simultaneously with an fMRI-compatible EEG equipped with standard Ag/AgCI electrodes from 64 sites on the scalp oriented in a 10-20 scheme system (Brain Products, Gilching, Germany). Initial fMRI pulse and ballistocardiographic artifact correction was completed in BrainAnalyzer 2 (Brain Products, Gilching, Germany) using classic subtraction and filtering approaches (Allen et al., 2000, 1998). These mid-level processed EEG measurements were then further processed using in-house software in MATLAB (Mathworks, Inc., Natick, MA, USA) and the EEGLAB toolbox (Delorme and Makeig, 2004; Mullen et al., 2013). Despite the subtraction and filtering approaches applied, residual artifact from the fMRI pulse persisted. To remove these lingering artifacts, we developed a new cleaning pipeline.

Our cleaning pipeline included steps tailored to remove common EEG artifact (e.g., eye blinks, muscle-related activity) and then targeted the high frequency noise in the 16-19 Hz and 34-38 Hz range. EEG data were bandpass filtered between 0.75 Hz and 50 Hz using a Finite Impulse Response (FIR) filter. Next, EEGLAB’s automated clean_rawdata function was used to determine channels that differed substantially from the estimated signal (derived from other channels) or had consistent flat-lining. Then, the EEG data were subjected to an Independent Component Analysis (ICA) decomposition and the ADJUST algorithm (Mognon et al., 2011) was used to remove ICA components associated with stereotyped noise. Following ICA decomposition, bad channels were interpolated using spherical interpolation. As a final step in EEG preprocessing, the EEG data were subjected to Artifact Subspace Reconstruction (ASR) (Chang et al., 2020; Mullen et al., 2015), which we used to target the aforementioned residual high frequency noise from the fMRI artifact. This method, in combination with the ICA cleaning method allows for the targeting of both stationary and non-stationary persistent artifacts. To deploy ASR on the dataset, we first created a “clean” reference signal from each subject’s EEG data by: 1) concatenating EEG segments that were at least 1000ms long with amplitude below 100μV, (2) and notch filtering (FIR) the EEG between 16-19 Hz and 34-38 Hz. Following the creation of the reference signal, ASR was then used to reconstruct the EEG that contained large fluctuations greater than 5 standard deviations beyond the reference signal (in 500ms chunks). Lastly, the data were re-referenced to a common average reference.

To construct EEG networks, the signal from each sensor was separated into standard frequency bands corresponding to δ (1-3Hz), θ (4-7Hz), α (8-13Hz), β (15-30Hz) and γ (30-60Hz) with a Butterworth filter (8^th^ order) followed by Hilbert transformation. Weighted functional connectivity adjacency matrices were constructed for each frequency band using the de-biased weighted phase-lag index (dwPLI) (Vinck et al., 2011). Each node in the adjacency matrix corresponds to a channel with the weight representing the strength (phase-lag) of the connection. Specifically, dwPLI is calculated as,

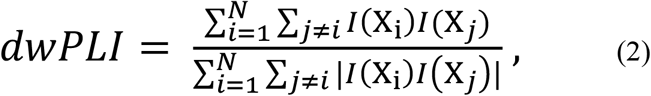

where *I(X_i_)* corresponds to the imaginary component of time series data (*X*) from channel *i*. Specifically, dwPLI is the sum of all pairwise products of the magnitudes of the imaginary components. In addition, dwPLI accounts for any bias due to the number of data points.

### 2.5 dMRI Acquisition and Preprocessing

Diffusion spectrum imaging (DSI) scans were acquired for each session. DSI scans sampled 258 directions using a Q5 half-shell acquisition scheme with a maximum b-value of 5,000 and an isotropic voxel size of 2.4 mm. Minimal preprocessing was carried out on the DSI scans and was restricted to motion correction. Following a similar procedure to the fMRI motion correction, motion was first assessed and applied for all of the b0 volumes, and a template was created for each scan composed of the average of the b0 volumes. Next, the b0 volumes and vectors were transformed using the estimated head motion correction, b0 template coregistration, b0 template-to-T1w coregistration and spatial normalization into MNI space using a single Hamming weighted sinc interpolation.

Fiber tracking was performed in DSI Studio (www.dsi-studio.labsolver.org) with an angular cutoff of 35°, step size of 1.0 mm, minimum length of 10 mm, spin density function smoothing of 0, and a maximum length of 250 mm. Deterministic fiber tracking was performed until 500,000 streamlines were reconstructed for each session. As with the fMRI volume data, streamline counts were estimated in 200 nodes using the same Schaefer 200 atlas (Schaefer et al., 2018) and 21 subcortical regions part of the Harvard-Oxford atlas (Makris et al., 2006). Connectivity matrices were then normalized by dividing the number of streamlines (*T*) between region *i* and *j*, by the combined volumes (*v*) of region *i* and *j*,

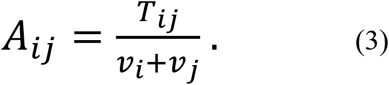

### 2.6 Graph Theoretical Analysis

We calculated nine commonly used and diverse graph metrics on each weighted dMRI, fMRI and EEG network. The graph metrics are: degree, clustering coefficient, characteristic path length, small-world propensity, global and local efficiency, synchronizability, spectral radius, and eigenvector centrality. See supplemental for detailed description of each network measure.

### 2.7 Degree

The weighted node degree (*k_i_*) is defined as the sum of all connections of a node (Rubinov and Sporns, 2010),

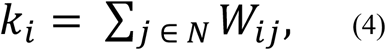

where *W* is the weighted adjacency matrix of a network with *N* nodes.

### 2.8 Clustering Coefficient

The weighted clustering coefficient (*C*) for node *i* is the intensity of triangles in a network (Onnela et al., 2005) and is calculated as,

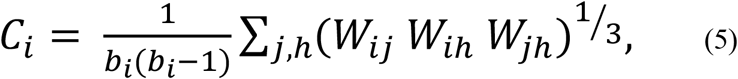

where *W* is the weighted adjacency matrix and *b* is the number of edges for node *i*.

### 2.9 Characteristic Path Length

The characteristic path length (*L*) is the average shortest path length between all nodes (Rubinov and Sporns, 2010),

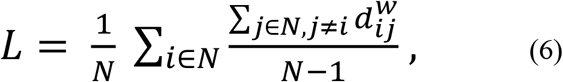

where 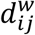 is the is the distance between nodes i and j. To calculate 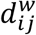, we first take the inverse of the edge weights to transform the weight to a measure of length (i.e., to transform a strong connection strength to a short length). We then determine the shortest path between nodes *i* and *j* (using the inverted weights), and 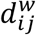 is the sum of the inverse of the edge weights along this shortest path.

### 2.10 Small-World Propensity

Small-world propensity (φ) quantifies the extent to which a network displays small-worldness, a network property that combines the presence of local clustering with a short path length, while factoring in variation in network density (Muldoon et al., 2016). Small-worldness is calculated as,

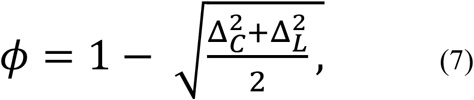

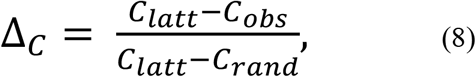

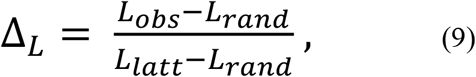

where *C_obs_* is the observed clustering coefficient and *L_obs_* is the observed characteristic path length of the network; *C_latt_, L_latt_, C_rand_*, and L_rand_ are clustering coefficient and characteristic path length from lattice and random networks with the same number of nodes and edge distribution.

### 2.11 Global and Local Efficiency

The efficiency of a node is the inverse of the path length (Rubinov and Sporns, 2010). Global efficiency (*E_g_*) is the inverse shortest path length,

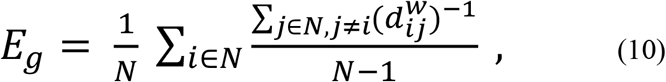

where 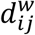 is the previously defined distance between node *i* and *j*.

Local efficiency (*E_l_*) is the global efficiency computed on the neighborhood of node *i*,

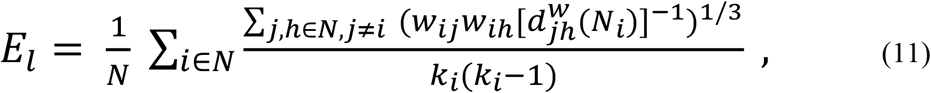

where *w_ij_* and *w_ih_* is strength of the connection between node *i* to *j* and *h*, respectively, and *d_jh_ (Ni)* is the length of the shortest path between nodes *j* and *h* that contains only neighbors of node *i*.

### 2.12 Synchronizability

Synchronizability is a measure of linear stability for a network of coupled dynamical systems (Motter et al., 2005),

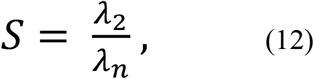

where *λ_2_* is the second smallest eigenvalue of the unnormalized Laplacian matrix (*L*) and *λ_n_* is its largest eigenvalue. The Laplacian is calculated as,

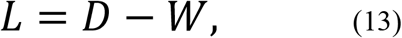

where *D* is the degree matrix of the weighted adjacency matrix, *W*.

### 2.13 Spectral Radius

The spectral radius measures the ease with which diffusion process can occur in a network. The spectral radius is calculated as,

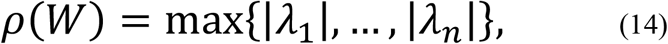

where |*λ*| corresponds to the absolute value of the eigenvalues of a network.

### 2.14 Eigenvector Centrality

Eigenvector centrality (*EC_i_*) measures how influential a node is in a network, with a high value indicating a node is connected to other highly influential nodes (Newman, 2008). The eigenvector centrality of node i is given by the i-th entry in the dominant eigenvector, which is the vector ***v***=[*v_1_*, … *v_N_*] that solves

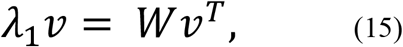

where *λ*_1_ is the largest eigenvalue of the weighted adjacency matrix, *W*.

### 2.15 Intra-class Correlation

The intra-class correlation (ICC) is a measure used to quantify the test-retest reliability of a measure. We used the ICC to measure the consistency of individual connections across the dMRI, fMRI and EEG networks and across the graph metrics for each network. To accomplish this, we calculated two variants of the ICC, the within (ICC_w_)- and between (ICC_b_)-subjects (Wei et al., 2004). ICC_w_ and ICC_b_ are, respectively, calculated as,

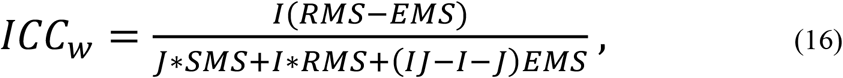

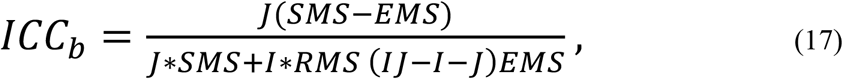

where *I* is the number of subjects and *J* is the number of sessions, *SMS, RMS and EMS* represent the ANOVA measures of mean square error between sessions, subjects, and due to error, respectively. The reliability of a measurement is considered: 1) “poor” if the ICC values is less than 0.4; 2) “fair” for ICC values between 0.4 and 0.6; 3) “good” for ICC values between 0.6 and 0.8; and 4) “excellent if ICC values exceed 0.8.

### 2.16 Fingerprinting Analysis

To perform a fingerprinting analysis, as in Finn *et al*., 2015, we quantified the degree of similarity between networks. This analysis was performed separately for each of the dMRI, fMRI and EEG modalities. Connectivity matrices were converted for each individual and run into a vector using the values from the upper triangle of the matrix resulting in vectors of 1 × 24,310 for dMRI and fMRI, and 1x 2,016 for EEG. Thus each vector, *p*, represents a single connectivity matrix for a given subject during a given session, and for functional matrices, in a given state (task/rest).

Next, separately within each modality, for each connectivity matrix (representing a subject, session, and state), we calculated the pairwise similarity between two vectors, *p* and *q*, using the Euclidian distance to create a dis-similarity matrix (*D*) where

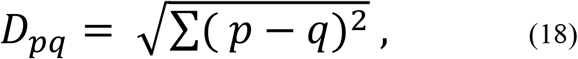

and each entry in *D_pq_*, corresponds to the dis-similarity between the brain network *p* to *q*. However, since the Euclidian distance formally assesses dis-similarity and we were interested in evaluating similarity, we converted from a dis-similarity to a similarity (*S*) measure by

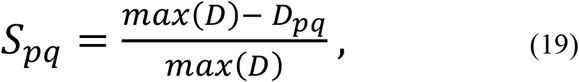

where *max(D)* corresponds to the largest value in matrix *D*. This normalization ensures that the similarity matrix *S* ∈ [0 1].

In order to perform a fingerprinting analysis, for each vector, *p*, we then looked for the entry *S_pq_* with the highest similarity value. If for this entry, the vectors *p* and *q* were from the same individual (but could be from different sessions or states), then the fingerprinting analysis was classified to be successful at identifying the individual.

Fingerprinting performance for each imaging modality was assessed using two measures. The first measure quantifies the overall fingerprinting accuracy across subjects, and was calculated as the percentage of matrices which were successful in identifying an individual. While this measure is useful from a classification standpoint, we were also interested in the level of separation between matrices within versus between individuals. Therefore, in the second measure, we assessed the separability (*T*) of each modality. The separability of each matrix, T*_p_*, was defined to be

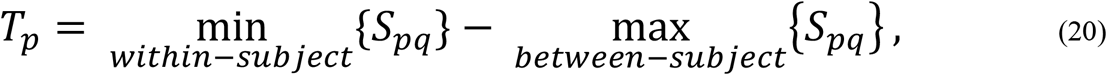

where the first term is constrained to *q* from the same subject as *p*, and the second term is constrained to q from all subjects other than p. The resulting values of *T* ∈ [-1 1], where a value of 1 indicates perfect similarity within a subject across sessions and no similarity to other subjects and, conversely, −1 indicates no similarity across runs within a subject.

### 2.17 Statistical Tests

Analysis of variance (ANOVA) was used to quantify the magnitude difference in ICC scores and the difference in the magnitude of the network similarity. Corresponding p-values were corrected for multiple comparison using Boneferroni correction. The Brain Connectivity Toolbox was used to calculate network measures (Rubinov and Sporns, 2010). All analyses were conducted in MATLAB 2017b.

## 3 Results

We analyzed the reproducibility of brain network properties derived from structural and functional brain imaging using the intra-class correlation (*ICC*). For the dMRI analysis, this involved analyzing brain networks from 25 subjects across 8 sessions for a total of 200 structural networks. For the fMRI and EEG analysis, a tradeoff between maximizing subjects and sessions was made across resting-state and tasks resulting in a range from 17-26 subjects, each with 6 sessions (see Methods section for details).

### 3.1 Reliability of Individual Connections

We first assessed the reliability of individual connections between brain regions or sensors. We calculated the ICC within a subject (ICC_w_) and between subjects (ICC_b_) for each connection across the three imaging modalities. As expected, we found that across imaging modalities, individual network connections are more reliable within-than between-subjects (Figure 1A and B). Across imaging modalities, individual edges exhibit high variability in their reliability scores, with ICC_w_ values ranging from poor (< 0.4) to excellent (> 0.8) reliability (Figure 1A). By contrast, ICC_b_ scores had consistently poor (< 0.2) reliability across all imaging modalities (Figure 1B). For dMRI, the mean ICC_w_ was 0.21 ± 0.24 (SD) and the mean ICC_b_ score was −1×10^−4^ ± 0.01 (SD). For resting-state fMRI the mean ICC_w_ was 0.23 + 0.13(SD). Lastly, for the EEG the α-band had the highest mean ICC_w_ (0.39 + 0.16(SD) compared to the other frequencies (δ: 0.03 ± 0.05(SD); θ: 0.09 ± 0.08(SD); α: 0.20 ± 0.12(SD); g: 0.10 ± 0.07(SD)). An ANOVA assessing differences across imaging modalities found significant differences in the ICC_w_ (F_6,85249_ = 2241; p_corrected_ << 0.001). One important feature is the long-tail distribution in the dMRI ICC_w_ indicating that a small number of connections have excellent (> 0.8) reliability. We additionally looked to see if there was a relationship between connection strength and reliability (Figure 1C-E).

**Figure 1.**
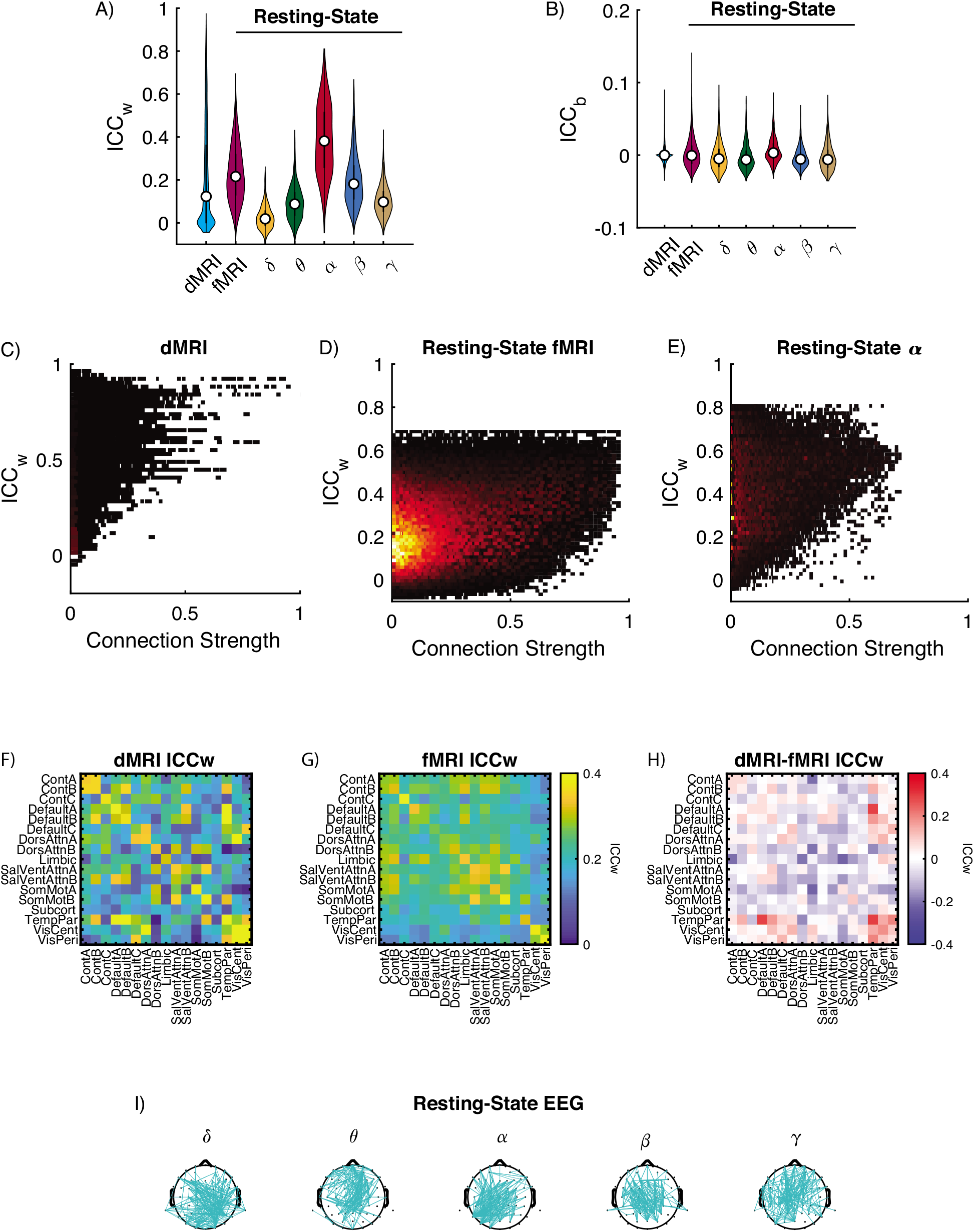
Reliability of individual connections. (A) Distribution of ICC_w_ and (B) ICC_b_ for dMRI and resting-state fMRI and EEG frequency bands. For each violin plot, the central dot indicates the median, and the line indicates the 25th to 75th percentiles. (C-E) Cumulative distribution plots showing the proportion of connections and corresponding connection strength from the top 10% (most reliable) and bottom 10% (least reliable) of ICC_w_ scores for (C) dMRI; Resting-State (D) fMRI and (E) EEG-α. F-H) Average reliability of connection within and between cognitive systems for (F) dMRI and (G) resting-state fMRI. H) Differences in average ICC_w_ scores across cognitive systems between dMRI minus the fMRI. (I) Connections with ICC_w_ scores in top 10% for δ, θ, α, β and γ frequency bands plotted on the scalp for resting-state EEG. Cognitive systems are defined as Cont: Control A/B/C, Default: Default Mode A/B/C, DorsAttn: Dorsal Attention A/B, Limbic, SalVentAtt: Salience/Ventral Attention A/B, SomMot: Somatomotor A/B, Subcortical, TempPar: Temporal Parietal, VisCent: Visual Central, VisPer: Visual Peripheral.

We next assessed if for dMRI and resting-state fMRI there is an association between ICC_w_ scores and cognitive systems. First, we mapped edgewise scores and then averaged over edges within each of the 17 cognitive systems from the Schaefer 200 layout combined with 21 subcortical regions from Harvard-Oxford atlas. As a trend, connections within a cognitive system for dMRI and resting-state fMRI exhibited the strongest reliability as can be seen from the figure because of the high values along the diagonal (Figure 1F and G, respectively). However, a direct comparison between dMRI and fMRI showed distinct distribution of reliability across cognitive systems. dMRI reliability was stronger within the Frontal-Parietal Control system and between the Visual, Default Mode, and Temporal Parietal systems (red entries in Figure 1H), while in fMRI, stronger values were distributed between cognitive systems (blue entries in Figure 1H). For the EEG data we could not perform the same mapping to cognitive systems, so instead resting-state ICC_w_ scores from the top 10% ICC_w_ distribution are plotted onto the scalp (Figure 1I).

Given the different cognitive demands associated with task performance, one might expect reliability scores during task states to differ from those at rest. However, when we examined task induced changes in reliability, we found that task associated ICC_w_ and ICC_b_ values for fMRI and EEG scores exhibited similar pattern to resting-state (Figures 2 and 3, respectively). To test for changes, we assessed an ICC x Task ANOVA and found that the ICC x Task interaction was significant (F*_10,501389_* = 1242, p_corrected_ << 0.001). For the EEG, we additionally added frequency as a variable in our ANOVA design and found that the ICC x Task x Frequency interaction was significant (F*_40, 201190_* = 140, p_corrected_ << 0.001) with the α-band having the highest ICC_w_ scores.

**Figure 2.**
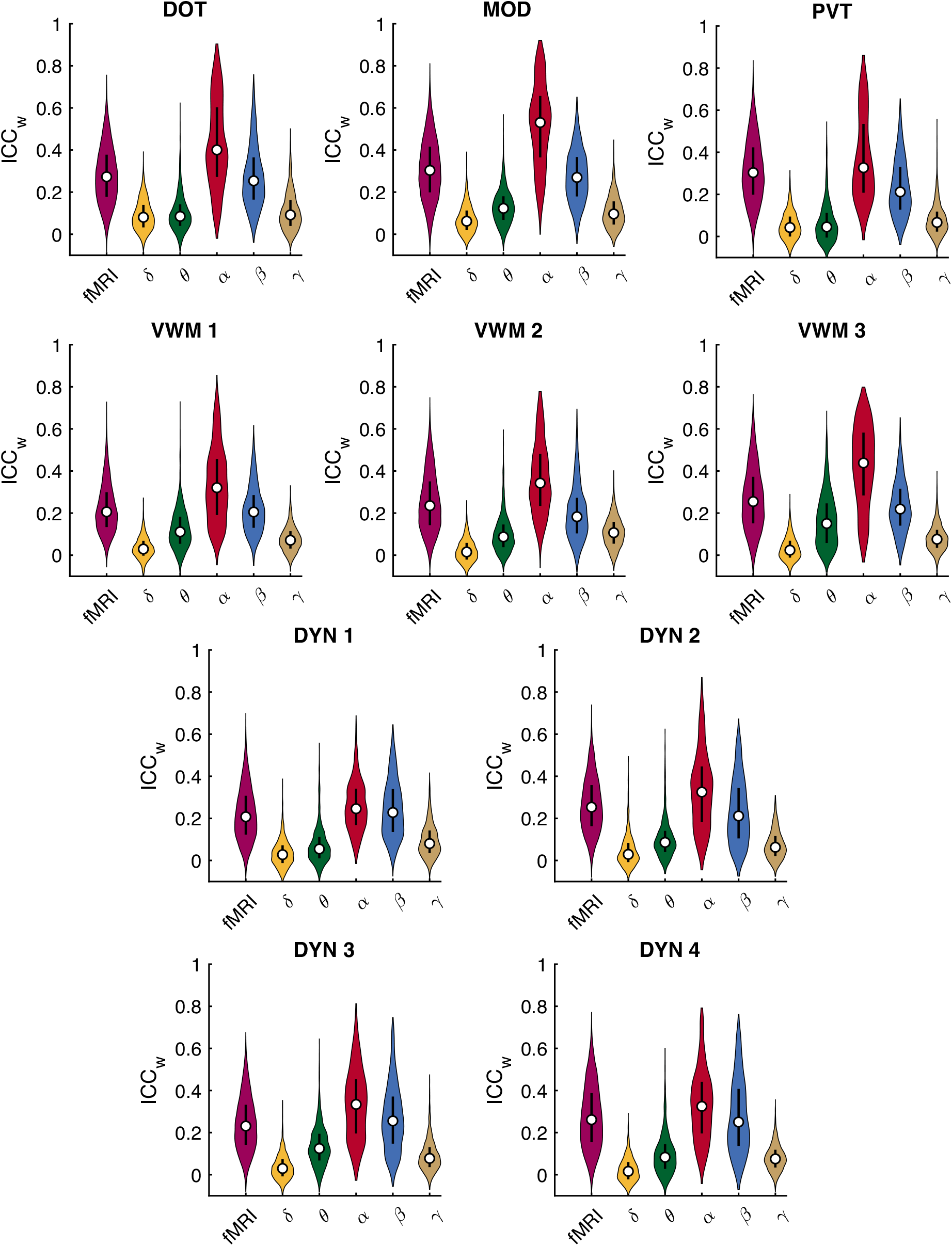
Reliability of individual connections within-subjects (ICC_w_) for fMRI and EEG frequency bands across tasks: DOT, PVT, MOD, VWM-1:3, and DYN-1:4. For each violin plot, the central dot indicates the median, and the line indicates the 25th to 75th percentiles. DOT: Dot Probe Task; DYN: Dynamic Attention Task; MOD: Modular Math Task; PVT: Psychomotor Vigilance Task; VWM 1-3: Visual Working Memory.

**Figure 3.**
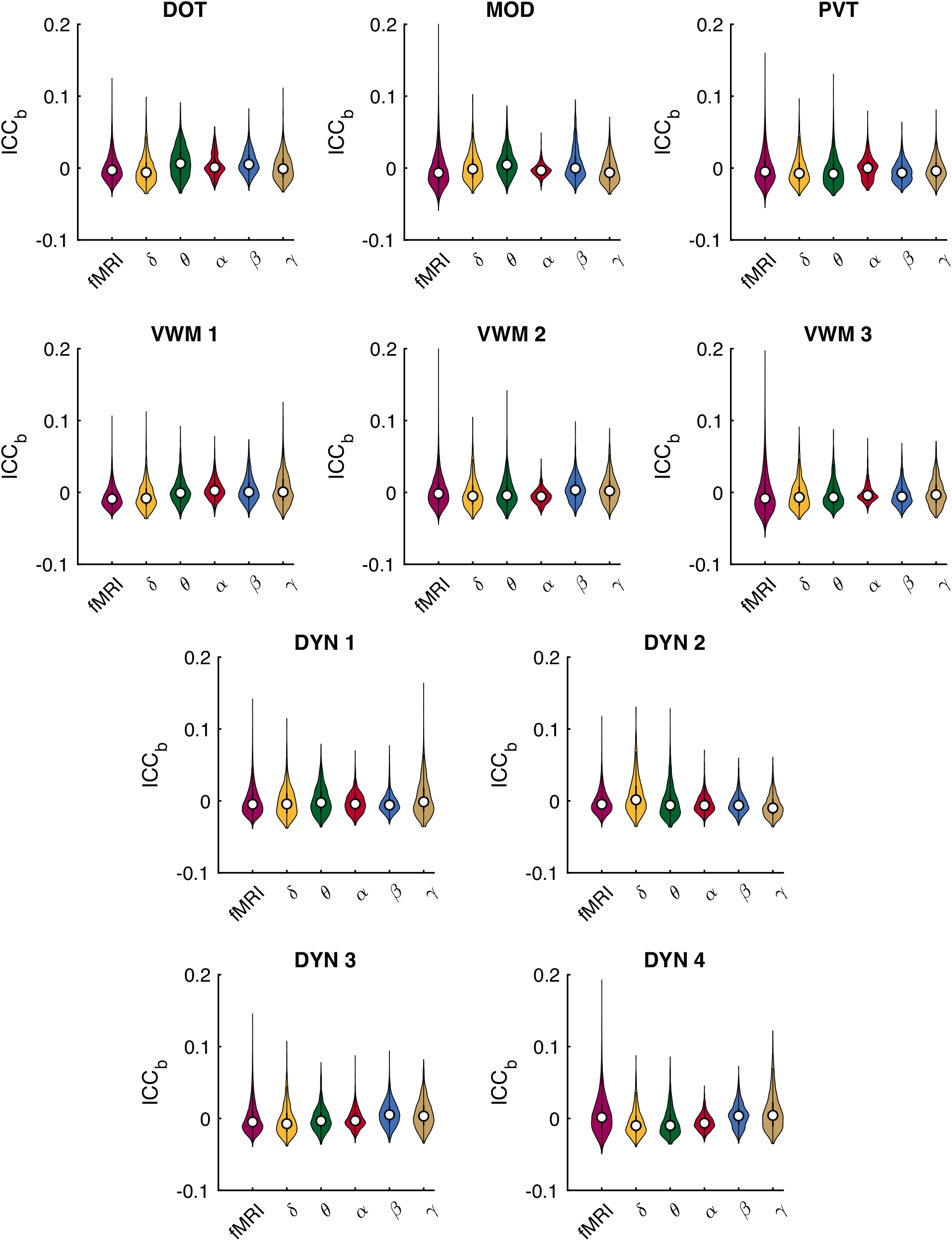
Reliability of individual connections between-subjects (ICC_b_) for fMRI and EEG frequency bands across tasks: DOT, PVT, MOD, VWM-1:3, and DYN-1:4. For each violin plot, the central dot indicates the median, and the line indicates the 25th to 75th percentiles. DOT: Dot Probe Task; DYN: Dynamic Attention Task; MOD: Modular Math Task; PVT: Psychomotor Vigilance Task; VWM 1-3: Visual Working Memory.

Similarly, we assessed if for task fMRI there is an association between ICC_w_ scores and cognitive systems. We mapped edgewise scores to the 17 cognitive systems in the same manner as for the resting-state and plotted the difference between the ICC_w_ values during task and resting-state in Figure 4. We generally observed higher reliability during task states, and found that for tasks with repeated sessions, the ICC_w_ progressively increased from resting-state as the sessions progressed (Figure 4 VWM and DYN tasks). For task EEG data, ICC_w_ scores from the top 10% of the ICC_w_ distribution were plotted onto the scalp and we did not notice any overt reconfiguration in scalp distribution from resting-state to task (Figure 5).

**Figure 4.**
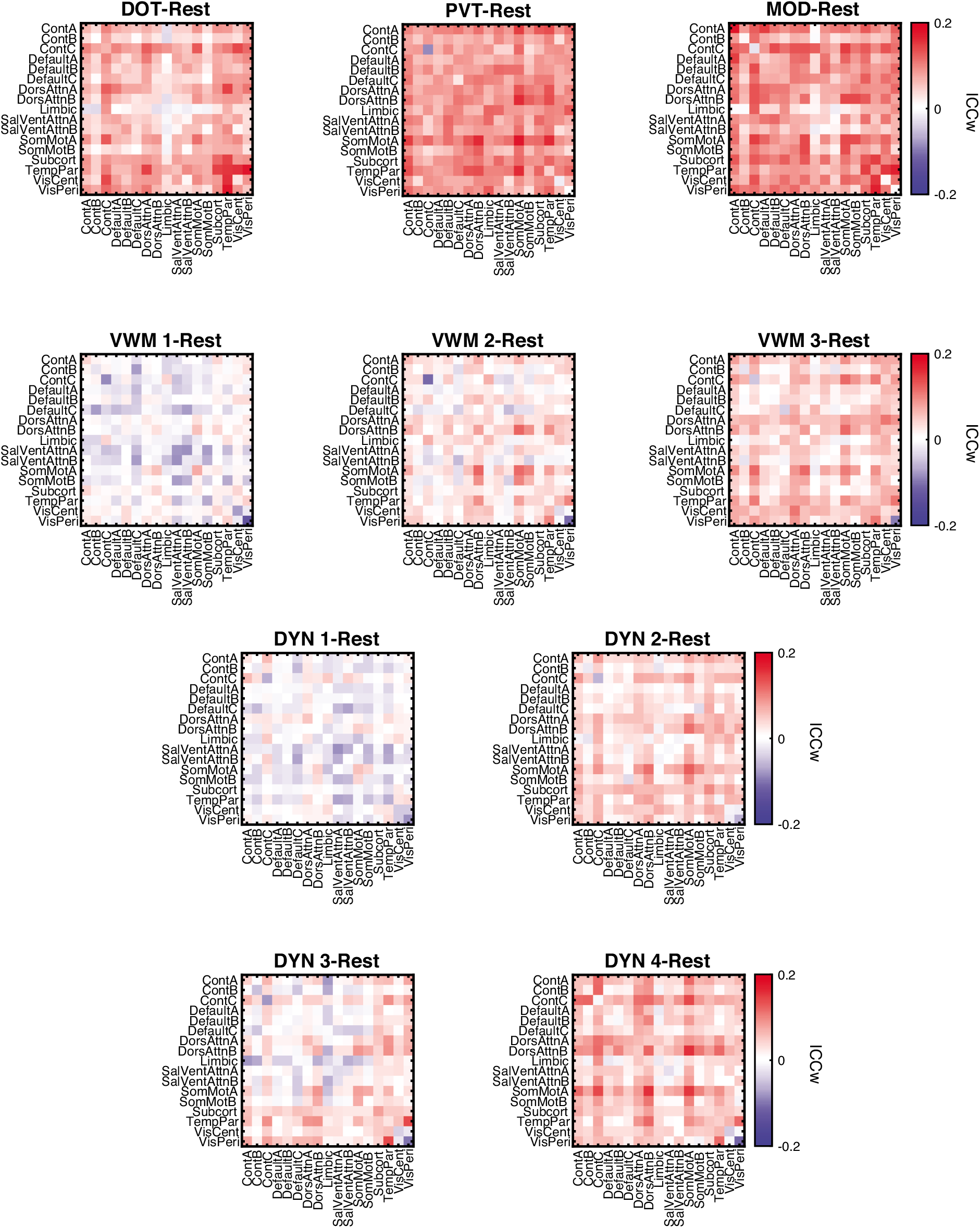
fMRI changes in reliability from resting-state for each task: DOT, PVT, MOD, VWM-1:3, and DYN-1:4. Connections are mapped unto 17 cognitive systems from Schaefer cortical and Harvard-Oxford subcortical atlas. DOT: Dot Probe Task; DYN: Dynamic Attention Task; MOD: Modular Math Task; PVT: Psychomotor Vigilance Task; VWM 1-3: Visual Working Memory.

**Figure 5.**
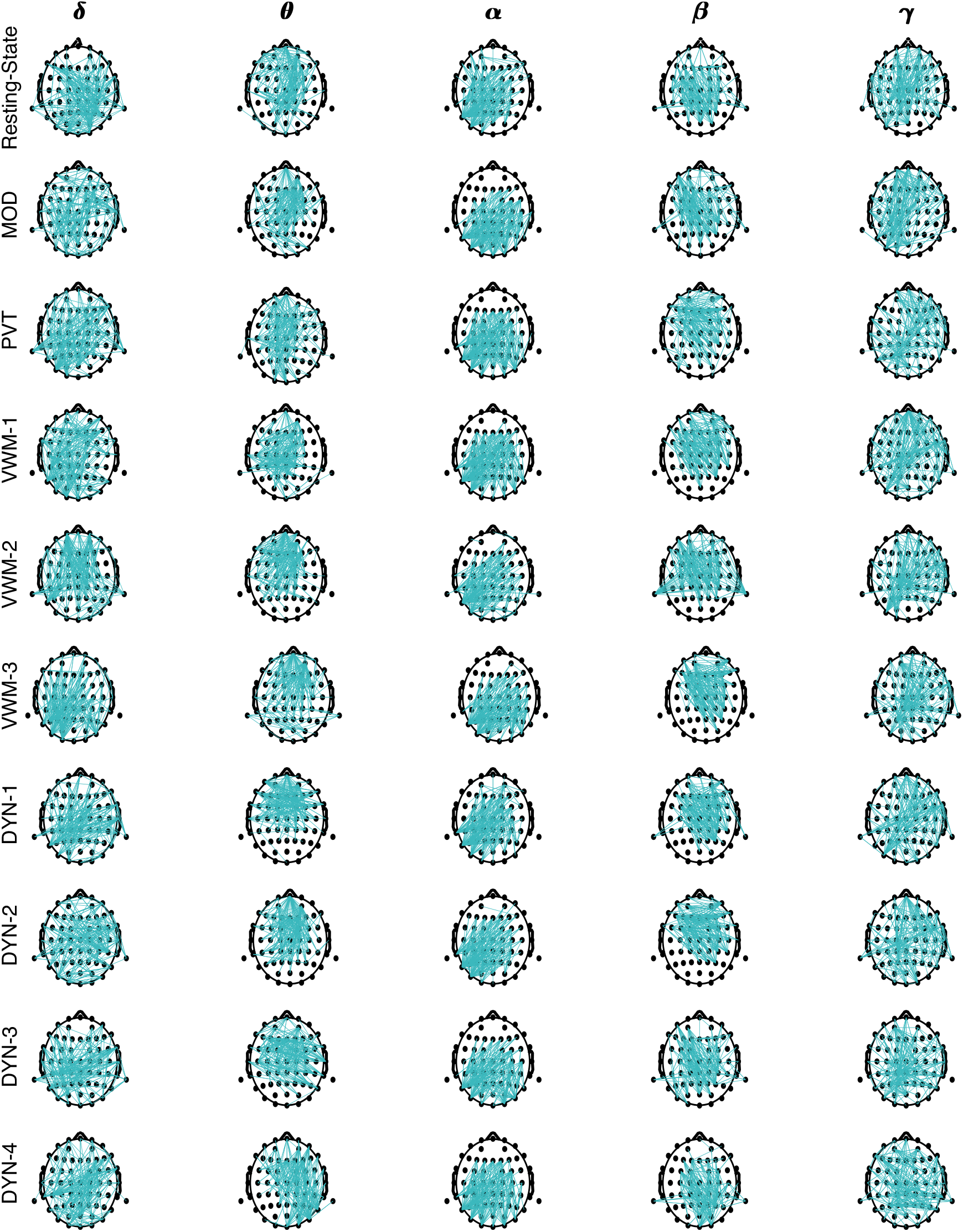
Scalp distribution across tasks for top 10% of ICC_w_ scores. For Resting-State and each task: DOT, PVT, MOD, VWM-1:3, and DYN-1:4, (10 in total), connections with ICC_w_ scores in top 10% for δ, θ, α, β and γ frequency bands are plotted on the scalp. DOT: Dot Probe Task; DYN: Dynamic Attention Task; MOD: Modular Math Task; PVT: Psychomotor Vigilance Task; VWM 1-3: Visual Working Memory.

### 3.2 Reliability of Network Measures

We next assessed the reliability of higher order network properties. For each brain network, nine measures were calculated along with their corresponding ICC_w_ and ICC_b_ scores. We found significant differences between modalities in ICC_w_ (F_6,66_ = 45; p-corrected << 0.001). As shown in Figure 6 A, across all imaging modalities and network properties, the dMRI exhibited the highest ICC_w_ scores (0.71 ± 0.06 (SD)). By comparison, resting-state fMRI exhibited relatively poor reproducibility (0.35 ± 0.12 (SD)), and EEG’s reproducibility was frequency dependent with the α-band having the highest ICC_w_ scores (0.43 ± 0.09 (SD)). ICC_b_ scores across all modalities were close to zero (Figure 6B).

**Figure 6.**
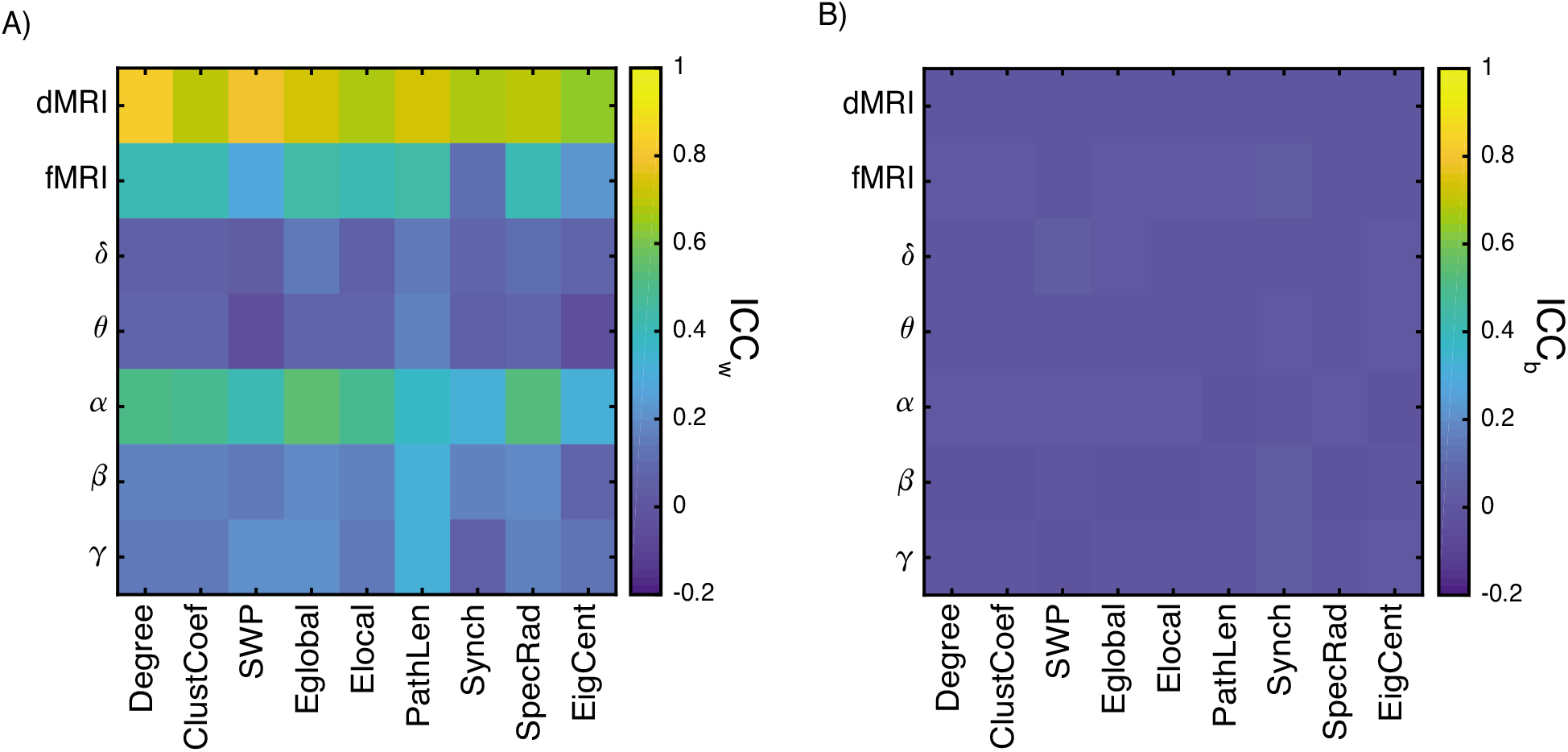
Graph Measures for dMRI and Resting-State fMRI and EEG for (A) ICC_w_ and (B) ICC_b_ values.

We next assessed if performing a task alters the reliability of network measures (Figure 7). For fMRI, we evaluated the Task x Network Measure x ICC ANOVA design and found significant interactions between Task x ICC (F*_8,80_* = 99, p_corrected_ << 0.001) and Network Measure x ICC (F*_10,80_* = 9.88, p_corrected_ << 0.001) (Figure 7A). For the EEG, we evaluated the Task x Frequency x Network Measure x ICC ANOVA design, and we found a significant interaction between Task x Frequency (F *_32,792_* = 5.35, p_corrected_ < 0.001) and Frequency x Network Measure (F*_40,792_* = 6.33, p_corrected_ < 0.001) (Figure 7A). From Figure 7 it is apparent that the α-band is the most consistent across resting- and task-state, while the β-band shows an increase in ICC_w_ in the task-states. It is also worth noting that Synchronizability and Eigenvector Centrality exhibited weaker ICC_w_ scores relative to the other metrics across resting- and task-states for both fMRI and EEG.

**Figure 7.**
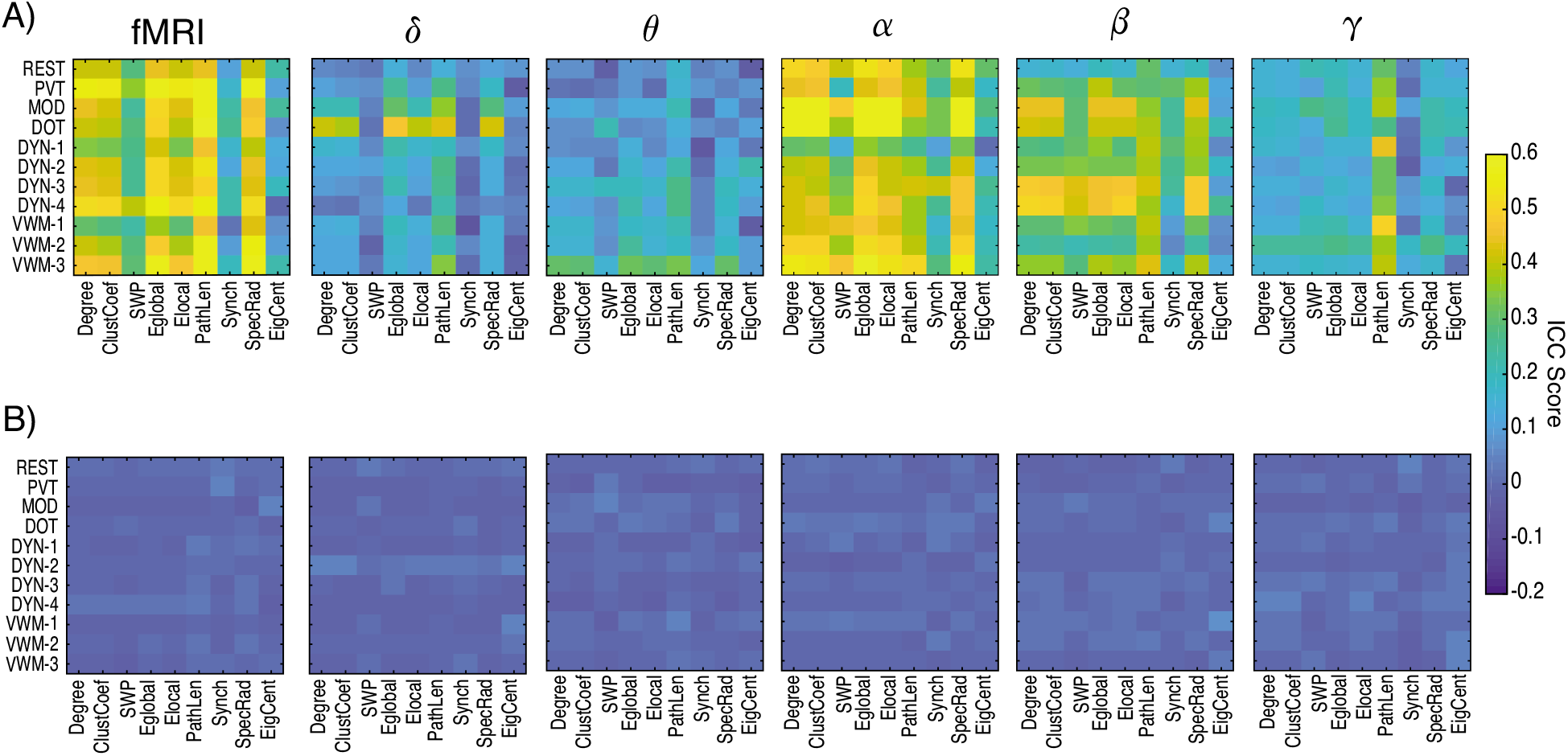
ICC values for network measures across task and resting-state. (A) ICC_w_ and (B) ICC_b_ values across tasks for fMRI and EEG frequency bands.

### 3.3 Fingerprinting Analysis

Our analysis so far has confirmed that dMRI networks are more reliable within a subject than fMRI and EEG networks. We therefore, expect that dMRI networks will have a higher probability of being able to identify an individual from a group, similar to a fingerprint (Finn et al., 2015). For functional networks, we would similarly expect the same of α-band EEG networks, given their relatively higher reliability scores. In order to fingerprint an individual, brain networks from the individual should be more similar to each other across runs relative to networks obtained from other individuals. To formally assess the similarity between brain networks, we measured similarity using the Euclidian distance (Methods). Our results indicate that fingerprinting was not uniform across all derived networks (F_6,168_ = 3402, p_corrected_ << 0.001). As expected, structural dMRI networks had the highest accuracy, but for functional networks, fMRI networks performed better than α-band EEG derived networks, and in fact, within EEG networks, β-band networks had the highest fingerprinting accuracy (Figure 8A).

**Figure 8.**
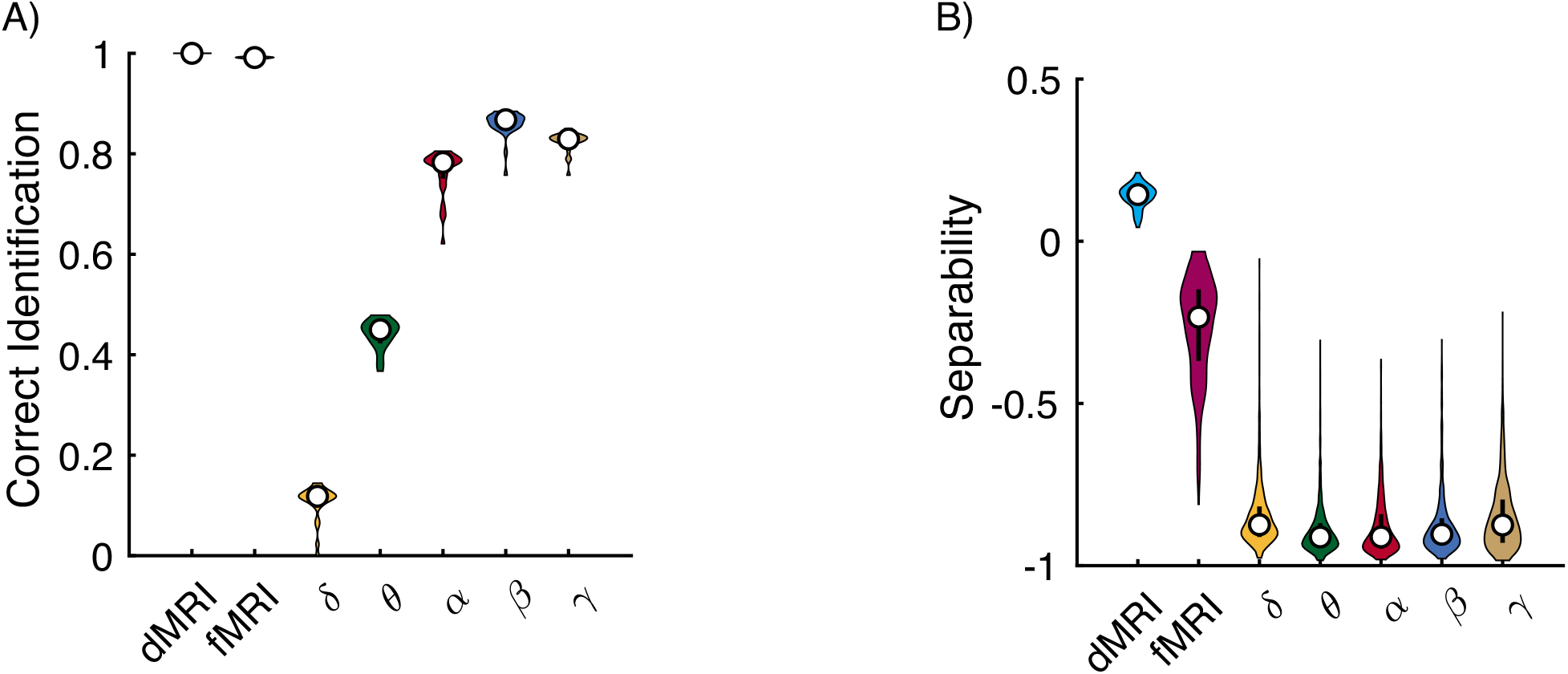
Fingerprinting performance across imaging modalities. (A) Proportion of networks that were correctly matched to the corresponding individual for dMRI, fMRI, and EEG derived brain networks. (B) Separability of each network in being matched to corresponding individual. For each violin plot, the central dot indicates the median, and the line indicates the 25th to 75th percentiles.

However, this analysis does not tell us about the separability across the networks derived from the different imaging modalities. Here we define separability as the difference in similarity between the minimum within-subject value for a network to the maximum between subject similarity for that network (Methods). Therefore, positive separability values indicate that a particular network for an individual is always more similar to other networks from that individual and negative values indicate the opposite. Separability values across imaging modalities were found to be significantly different (F_6,10093_ = 8618; p_corrected_ << 0.001). In addition, despite dMRI and fMRI having similar accuracy in fingerprinting, dMRI networks were more separable than fMRI and EEG (dMRI: 0.14 ± 0.04 (SD); fMRI: −0.26 ± 0.27 (SD); δ, θ, a, α and g: < −0.85 (mean)) (Figure 8B).

## 4 Discussion

In the current work, we analyzed the reproducibility of multimodal and multi-task structural and functional brain networks in a unique longitudinal and multi-modal data set with simultaneous EEG-fMRI recordings. In our analysis, each subject contained brain networks derived from dMRI, fMRI and EEG data, allowing us to assess how reliability differed in brain networks derived from different modalities and across task states.

### 4.1 Edgewise Reliability Differences Between dMRI, fMRI, and EEG

We first assessed the reliability of individual connections in the structural and functional brain networks and found stronger within-than between-subject reliability across all imaging modalities, in line with previous results (Birn et al., 2013; Noble et al., 2019, 2017; O’Connor et al., 2017; Pannunzi et al., 2017; Shehzad et al., 2009). The most reliable connections were also the ones that tended to be the strongest, corroborating previous findings in fMRI networks (Noble et al., 2017; Pannunzi et al., 2017). In addition, these connections when mapped on to cognitive systems, exhibited distinct patterning. As a trend, for dMRI and resting-state fMRI, connections within a cognitive system exhibited the strongest reliability, consistent with previous studies in functional networks (Birn et al., 2013; Noble et al., 2017; O’Connor et al., 2017; Shehzad et al., 2009). However, a direct comparison between dMRI and resting-state fMRI showed distinct distribution of reliability across cognitive systems. dMRI reliability was strongest within the Frontal-Parietal Control system and between the Visual to Default Mode and Temporal Parietal system, while in resting-state fMRI stronger values were distributed between cognitive systems.

When assessing task mediated changes, we found an increase in reliability across most tasks relative to resting-state in fMRI networks. In addition, we observed an increase in this reliability across multiple sessions of a given task, potentially indicative of an effect of learning the task. This finding compliments results from a previous study that found adding task-state fMRI networks improves predictive outcomes relative to resting-states fMRI (Gao et al., 2019).

For EEG, the α- and β-bands had the highest reliability scores for both resting- and task-states, confirming previous results (Kuntzelman and Miskovic, 2017). The strong reliability for the α- and β-band could be due to the fact that these frequencies are consistently activity, while the other frequency bands tend to have transient activity. In a similar manner to fMRI, EEG reliability increased during a task, but this increase was primarily in the α- and β-bands. In addition, we found no major changes when we mapped connections on the scalp from resting-state to task-state. This could be due to the low spatial resolution of EEG (Nunez et al., 1997).

### 4.2 Reliability of Graph Theoretical Measures

When examining the reliability of higher order network properties, we found that network properties had overall stronger reliability scores than individual connections in line with previous findings of Braun et al., 2012. This might lead one to ask how the prevalence of low reliability scores across most connections could produce fair to excellent reliability in higher order network properties? This result could be due to the fact that edges with higher reliability scores are associated with the stronger connections. Our graph theoretical properties are dependent on connection strength, and the stronger the connection, the more variance it accounts for in the higher order network values. Thus, despite most connections having poor reliability, the few strong connections with good to excellent reproducibility have a disproportionately higher impact on the reliability of a network measure. The notable exception is that in fMRI and EEG, synchronizability and eigenvector centrality had lower reliability scores than the other network properties. One possible reason for this is that these measures, particularly eigenvector centrality, are very sensitive to the state of the subject (Lohmann et al., 2010). These results indicate these measures might be more sensitive to detecting meaningful differences between individuals in studies where one is attempting to link functional brain connectivity to task performance or behavior.

We also found task associated differences in reliability for the fMRI and EEG. However for the EEG, the strongest increases in reliability were in the α- and β-bands. However, in contrast to Deuker *et al*., 2009 we did not find a corresponding increase in ICC_w_ scores in the δ and θ bands with task.

### 4.3 Fingerprinting

We found that dMRI and fMRI outperformed EEG derived networks in fingerprinting. However, the separability was not equal across these networks, with dMRI outperforming all functional networks. This is likely due to the fact that, unlike functional connectivity, structural connectivity is not state dependent.

It has been found that brain activity measured with fMRI is stable over time (Braga and Buckner, 2017; Gratton et al., 2018; Horien et al., 2019; Laumann et al., 2015) and in fMRI, within-subject variance can be reduced with high quality data with long scan times (~15 minutes) and multiple sessions (Birn et al., 2013; Laumann et al., 2015; Noble et al., 2017; Pannunzi et al., 2017). It has been argued that large amounts of data are needed in order to differentiate between true and artifact induced variance (Gordon et al., 2017; Power et al., 2012) and previous studies have found that reliability increases with more data (Anderson et al., 2011; Birn et al., 2013; Laumann et al., 2015; Noble et al., 2017; Shou et al., 2013). This high quality data is important because Horien *et al*., 2019 found that motion characteristics can be unique to an individual and can fingerprint a subject at a level greater than chance. In our data, individual scan times were limited to approximately 5 minutes, but data was collected over multiple sessions for a relatively large number of subjects, suggesting that we might expect more reliable results. However, our observation of the relatively weak accuracy and separability of EEG (a more direct measure of neuronal activity than fMRI) in fingerprinting an individual raises questions as to whether the increase in fingerprinting performance in fMRI on long time scans is based on neuronal activity. Also, respiration induced artifacts in fMRI exhibit the same stability over time (Power et al., 2019), which could also lead to increased reliability measurements.

Our direct comparison of fingerprinting between structural and functional networks indicates that structural networks more sensitive. In addition, these results indicate that the patterning in structural connectivity is far more unique to an individual than those in corresponding functional networks. These results suggest that structural networks might have more discriminative power than functional networks.

Unique brain connectivity features have previously been proposed to play a role in differences underlying behavior and cognition (Kanai and Rees, 2011). Specifically, difference in behavioral performance in motor and decision associated tasks are correlated with fractional anisotropy of the corpus callosum (Johansen-Berg et al., 2007; Westerhausen et al., 2006), optic radiation (Tuch et al., 2005) and grey matter density (Van Gaal et al., 2011). Cortical thickness within the superior parietal lobes has been found to be correlated with the rate of switching in a perception based task (Kanai et al., 2010). In addition structural features unique to an individual lead to characteristic brain functional activity in modeling analysis and task performance (Bansal et al., 2019, 2018a).

### 4.4 On Reliability, Confounding Variables, and Utility

Is a connection with poor reliability good or bad? To answer this, we need to be mindful of the goal at hand. First and foremost, we need to make sure that reliability values are not due to noise in the signal. On the other hand, if we are confident that low reliability is a genuine part of the signal, then that is also a very informative finding. The seminal work of Poldrack et al., 2015 found that functional connectivity exhibits a high level of variability within the same person over the course of a year. Along these lines, Noble et al., 2017 found that functional connections with strong reliability are not very informative when it pertains to predicting behavior. However, we need to be mindful that this is an effect limited to functional connectivity. Therefore, structural connections and/or higher-order network metrics might exhibit a stronger association between reliability and behavior. Also, finding highly reproducible brain connections and/or measures might be very important if we are looking for deviations from expected values that could be used as biomarkers for disease identification/progression. Alternatively, connections and/or measures with low reliability might be useful for studying individual differences and making correlations between structure and performance/behavior.

But, even beyond reliability and noise, our functional results could, along with previous literature, reflect the natural day-to-day changes in our brain. Neuroplastic changes in the brain are the hallmark of learning and memory (Lamprecht and LeDoux, 2004), and these changes or natural fluctuations and modifications in the *neural code* (Fairhall et al., 2001), reflecting learning and memory could be reflected in functional connectivity. Indeed, there are many examples of rapid neuroplastic changes in the brain that results in functional connectivity changes (e.g., Nierhaus et al., 2019), but see Perich et al., 2018 as an alternative theory. Moreover, in this particular dataset, individuals were recruited to capture substantial variability in sleep without experimental manipulation. While there is a substantial literature on brain related decrements due to sleep deprivation (Boonstra et al., 2007; Hudson et al., 2020) little is known about naturalistic fluctuations in sleep (Moturu et al., 2011; Thurman et al., 2018). These individuals, instead could be more “plastic” (or “stationary”) than other individuals. Future studies may disentangle these alternatives from a *reliability* explanation of our results.

fMRI-based analysis has been around for over two decades, but its clinical use has been limited, raising questions about its usefulness as a diagnostic tool. In addition, given that the effectiveness of any diagnostic tool is only as useful as it can be applied to an individual, then in this regard, structural networks should take a more prominent role in medicine. Regardless, one must consider how measures of reliability relate to the modality being studied, the state of the brain, and the question at hand in order to meaningfully ask questions about how brain networks change with disease or how individual differences in structure relate to performance and behavior.

## Acknowledgements

Funding: This research was supported by mission funding to the US CCDC Army Research Laboratory as well as sponsored by the Army Research Office and accomplished under Cooperative Agreement Numbers W911NF-10-D-0022 and W911NF-10-D-0002. Additional support was provided through University at Buffalo startup funding. The views and conclusions contained in this document are those of the authors and should not be interpreted as representing the official policies, either expressed or implied, of the Army Research Laboratory or the US Government.

